# Synchronized eye movements predict test scores in online video education

**DOI:** 10.1101/809558

**Authors:** Jens Madsen, Sara U. Julio, Pawel J. Gucik, Richard Steinberg, Lucas C. Parra

## Abstract

Experienced teachers pay close attention to their students, adjusting their teaching when students seem lost. This dynamic interaction is missing in online education. We propose to measure attention to online videos remotely by tracking eye movements, as we hypothesize that attentive students follow videos similarly with their eyes. Here we show that inter-subject correlation of eye-movements during instructional video presentation is substantially higher for attentive students, and that synchronized eye movement are predictive of individual test scores on the material presented in the video. These findings replicate for videos in a variety of production styles, learning scenarios and for recall and comprehension questions alike. We reproduce the result using standard web cameras to capture eye-movements in a classroom setting, and with over 1,000 participants at-home without the need to transmit user data. Our results suggest that online education could be made adaptive to a student’s level of attention in real-time.

## Introduction

In a classroom the level of attention is quite variable [1, 2]. An overt indicator of attention is the point of gaze [3, 4]. When students are not following the relevant teaching material, there is a good chance that they are not paying attention and that they will perform poorly in subsequent exams. Experienced teachers know this and adjust the interaction with students accordingly [5]. During online education this immediate feedback is lost. Here we suggest that standard web cameras could be used to monitor attention during online instruction based on the student’s eye movements.

In the context of online media eye tracking has been used extensively to evaluate user interfaces, advertising or educational material [6, 7]. In education research eye tracking has been used to improve instructional design, determine level of learner expertise, or purposefully guide eye movements during instruction [8]. These studies often focus on the content of eye fixations in static media, to determine, for example, whether users look at a specific graphic or whether they read a relevant text [9, 10]. This approach requires detailed analysis and interpretation of the specific content, and cannot be used routinely to evaluate individual students. Evaluating the content of eye fixations is particularly complicated for dynamic stimuli such as instructional video, which is increasingly abundant online. Here we focus on dynamic video and whether students “follow” that dynamic content, in the literal sense of following with their eyes.

Previous studies have shown that eye movements are correlated across subjects during video presentation [11, 12]. This inter-subject correlation (ISC) of eye-movements is elevated for dynamic, well-produced movies and video advertising [13–15], and is affected by the viewing task [16]. A variety of eye tracking measures have been used in educational research [17]. However, the observation that eye movements are synchronized across subjects has not been widely explored in the context of education. In particular, it has not been established yet whether inter-subject correlation of eye-movement depends on attention, or whether it is predictive of learning. Much of our eye movements during video seems to be driven by the visual dynamic [13], resulting in similar scan paths even when movies are presented backwards in time [11]. Thus, a remarkable fraction of eye movements seems to be guided by “bottom-up” processing of salient visual events in the video [18].

We hypothesize that typical online instructional videos synchronize eye movements across students, however, the level of synchrony depends on whether students are paying attention. Therefore, the correlation of eye-movement between subjects should be predictive of retention of the material presented in the video. The alternative hypothesis is that the stimuli drive eye movements without engaging a student’s mind meaningfully in the material. One may also argue that static stimuli, while not reliably guiding eye movements, may nonetheless engage students minds [19, 20].

We test this hypothesis by measuring ISC of eye movements and pupil size, recorded while a group of students watch short informal instructional videos typically found online. Consistent with the hypothesis, we find that significant inter-subject correlation, which drops in magnitude when viewers are distracted by a secondary task. Additionally, ISC of individual students with the group is predictive of individual performance in a subsequent test of recall and comprehension. To determine the robustness of these findings, we repeat the experiment for different learning scenarios and instructional videos produced in different styles. Finally, we replicate the results with remote students, using subjects own computers to capture their eye movements, without the need to transfer data from the user, thus preserving online privacy.

## Results

### Effects of attention on eye movements during video presentation

To test the hypothesis that synchronization of eye movements depends on attentional state, we recruited 60 subjects to participate in a series of experiments where they were asked to watch 5 or 6 short instructional videos in the laboratory while we monitored their eye movements. The videos covered a variety of topics related to physics, biology and computer science (Tab. 1). The videos reflected the most common contemporary formats, which feature a teacher writing on a board, or more modern storytelling using animations or the popular writing-hand style. A first cohort of subjects (N=27, 17 females, age 18-53 mean=26.74, standard deviation SD=8.98) watched 5 short instructional videos, after each video they took a test with questions related to the material presented in the videos. In the first cohort, subjects were told to expect this subsequent test. After watching the videos and answering questions they watched the videos again. To test for attentional modulation of ISC, in the second viewing subjects performed a serial subtraction task (count silently in their mind backwards in steps of seven starting from a random prime number between 800 and 1000). This is a common distraction task in visual attention experiments [21]. During the first attentive viewing eye movement of most subjects are well correlated (Fig. 1a). As predicted, during the second, distracted viewing eye movements often diverge (Fig. 1b). The same appears to be true for the fluctuations of pupil size. To quantify this, we measure the Pearson’s correlation of these time courses between subjects. For each student we obtain an inter-subject correlation (ISC) value as the average correlation of that subject with all other subjects in the group. We further average over the three measures taken, namely, vertical and horizontal gaze position as well as pupil size. This ISC is substantial during the normal viewing condition (Fig. 1c; ISC median=0.32, interquartile range IQR=0.12, across videos) and decreases in the second distracted viewing (ISC median=0.11, IQR=0.07). Specifically, a three-way repeated measures ANOVA shows a very strong fixed effect of the attention condition (*F*(1, 231)=749.06, *p*=1.93·10^−74^) a fixed effect of video (*F*(4, 231)=32.29, *p*=2.23·10^−21^) and a random effect of subject (*F*(26, 231)=9.21, *p*=1.62·10^−23^). For a replication of these results on 2 other experiments see Supplement S1. This confirms the evident variability across films and subjects. The predicted effect of attention, however, is so strong that despite the variability between subjects one can still determine the attention condition from the ISC of individual subjects (Fig. 1c). Specifically, by thresholding on ISC one can determine with high accuracy the attentional state of the subject (area under the receiver-operator curve of *Az* = 0.944± 0.033 – mean ± SD over videos). To determine which time scale of the eye dynamic dominates this inter-subject correlation we computed ISC resolved by frequency (Fig. 1d). We find that ISC and its modulation with attention are dominant in a time scale of 0.12-30 seconds for eye movements (0.03-8Hz) and 0.4-30 seconds for pupil size (0.03-2.6Hz).

**Figure 1:**
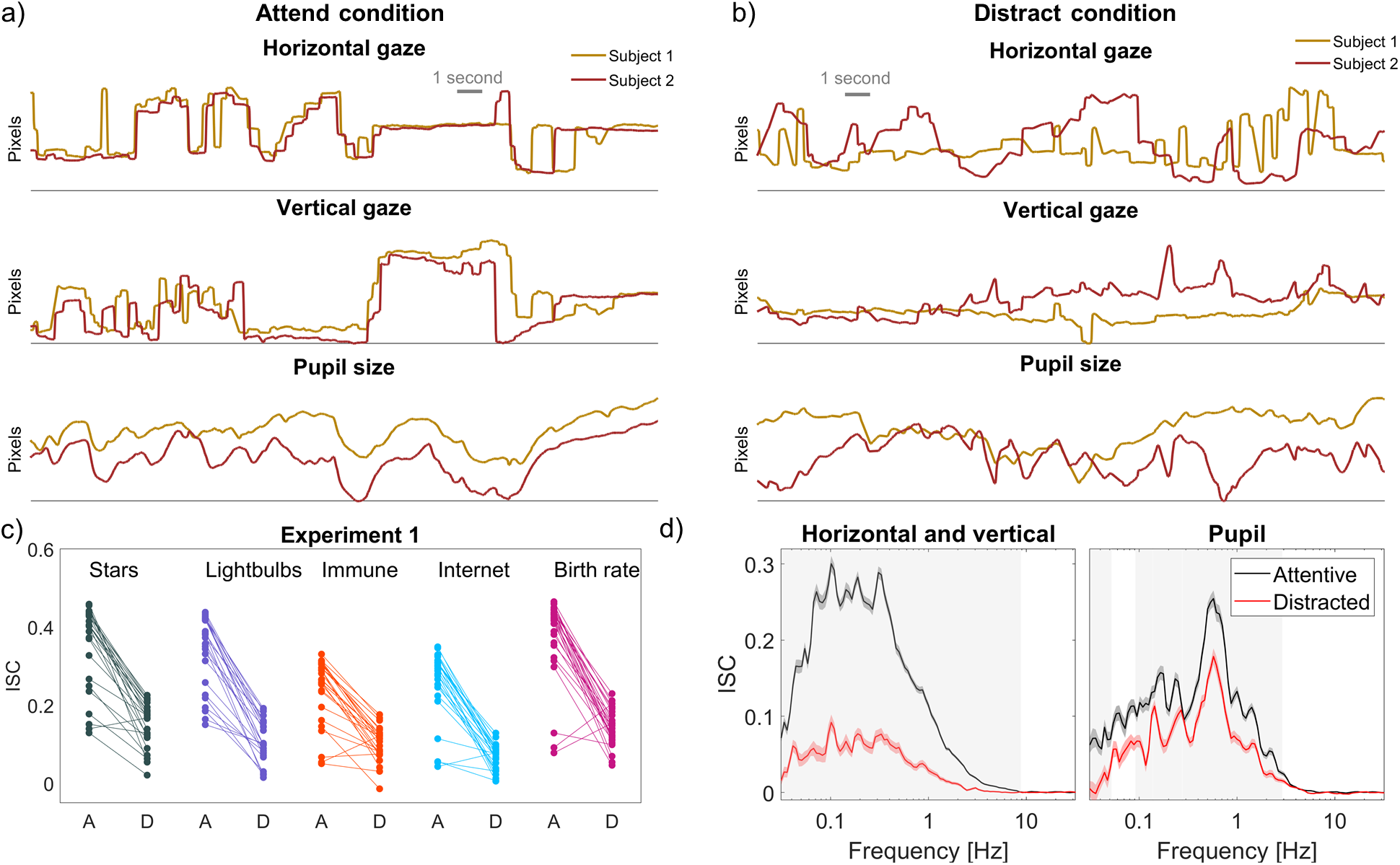
Inter-subject correlation of eye movements modulated by attention when watching instructional videos. **a)** Two subjects’ gaze position and pupil size follow each other during attentive viewing. **b)** The same two subjects viewing the same segment of video while distracted by a counting task. **c)** For each subject, inter-subject correlation (ISC) is measured as the mean correlation of vertical and horizontal gaze position and pupil size with that of other subjects. Values for each subject are shown as dots for all videos in Experiment 1. Each dot is connected with a line between two different conditions, namely, when subjects were either attending (A) or were distracted (D) while watching the video. **d)** ISC for the attentive and distracted conditions resolved by frequency, i.e. computed on band-pass filtered eye movements and pupil size. Each ISC value is averaged over the 5 videos and all subjects. In both panels significant differences between attending and distracted conditions are established using a cluster permutation test (grey shaded area indicates p<0.01).

**Table 1:**
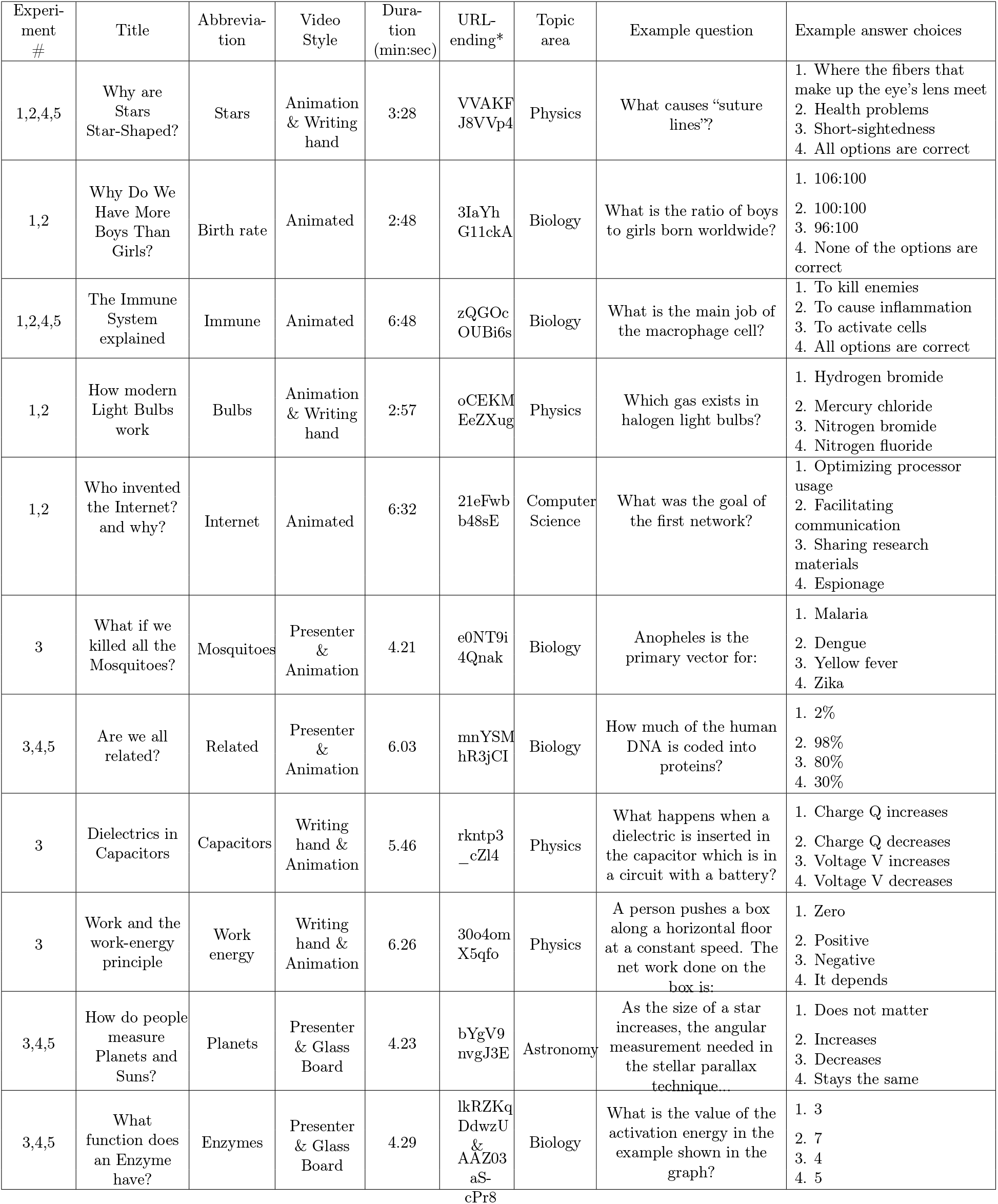
Experiment, title, abbreviation, style, duration, web address, and example questions and answer choices. Full list of of questions and answer options can be found here. URL of videos beginning with https://www.youtube.com/watch?v=

### Correlated eye movements and pupil size as predictors of test scores

In the previous experiments we confirmed the hypothesis that if subjects are distracted the ISC of eye movements and pupil size is reduced. Given the well-established link between attention and memory we therefore expect that ISC will be predictive of how much each subject retained from the instructional video. We tested this hypothesis by quizzing subjects after they had watched the video using a short, four-alternative forced-choice questionnaire (11-12 questions per video). Students that watched the video performed significantly better than naïve students (65.2% ± 18.8% versus naïve: 45%±8.8%; t(56)=-5.37 *p*=1.58·10^−6^; see Methods section for details). Importantly, we find a strong correlation between ISC and test scores across subjects for all videos we tested (Fig. 1b; r=0.61 ± 0.06, SD across 5 videos, *p*<3.60·10^−3^). This is true for ISC of eye movement and pupil size alike, even when luminance fluctuations are regressed out from the pupillary response (Supplement Fig. S6). Evidently, subjects with lower ISC performed poorer on the tests (e.g. subject 3 in Fig. 2a). Inversely, subjects with more correlated eye movements obtain higher test scores (e.g. subject 1 & 2 in Fig. 2a). This suggests that subjects who did not follow the visual dynamics of the video with their eyes were not paying attention and as a result their test scores were lower (see Supplement S1 for replication of these results on 2 other experiments). A statistical model of the data favours this causal interpretation (Supplement S3). However, the present study is only observational and the source of the correlation observed here between ISC and test performance remains undetermined.

**Figure 2:**
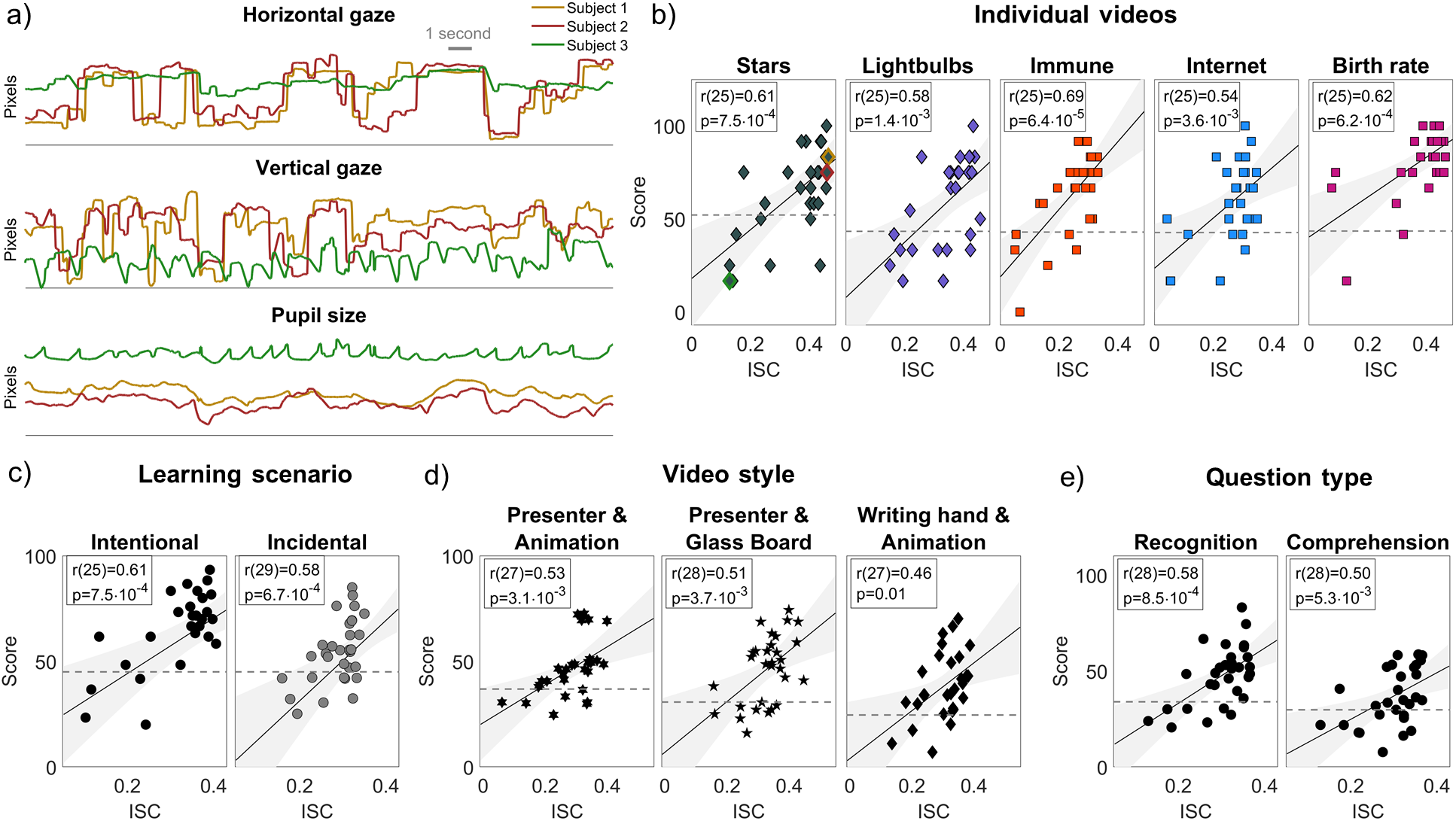
Inter-subject correlation of eye movements during instructional videos predicts learning performance. **a)** Eye movements of three representative subjects as they watch “Why are Stars Star-Shaped?”. Two high performing subjects have similar eye movements and pupil size dynamic. A third, low performing student does not match their gaze position or pupil size. **b)** Inter-subject correlation of eye movements (ISC) and performance on test taking (Score) for each of five videos in Experiment 1. Each dot is a subject. The high and low performing subjects (subjects 1-3) from panel (a) are highlighted for the Stars video. Dotted lines represent performance of subjects naïve to the video. **c)** Same as panel (b) but averaging over the 5 videos. The data was collected in two different conditions: During intentional learning (Experiment 1) where subjects knew they would be quizzed on the material. During incidental learning (Experiment 2) where subjects did not know that quizzes would follow the viewing. **d)** Videos in three different production styles (Experiment 3) show similar correlation values between test scores and ISC. Each point is a subject where values are averaged over two videos presented in each of the three styles. (See Supplement Fig. S2 for results on all 6 videos.) **e)** A similar effect is observed for different question types. Here each point is a subject with test scores averaged over all questions about factual information (recognition) versus questions requiring comprehension. ISC were averaged over all 6 videos in Experiment 3.

### Different learning scenarios

To test for the robustness of the effect we repeated the experiment, but this time subjects did not know that they would be quizzed on the content of the videos (Fig. 2c). The two scenarios thus constitute intentional and incidental learning which are known to elicit different levels of motivation [22]. As expected, we find a higher ISC in the intentional learning condition (ISC median=0.325, IQR=0.12, N=27) as compared to the incidental learning condition (ISC median=0.317, IQR=0.06, N=30) (two-tailed Wilcoxon rank sum test: z=2.67, *p*=7.68·10^−3^). This suggests that lower motivation in the incidental learning condition resulted in lower attentional levels and thus somewhat less correlated eye movements and pupil size. In the intentional learning condition test scores where higher as compared to the incidental learning condition (intentional learning score=65.22 ± 18.75 points, N=27, incidental learning score = 54.53 ± 15.31 points, N=31; two-sample t-test: t(56)=2.39, *p*=0.02, d=0.63). This may reflect increase motivation or more simply the increased difficulty of having to answer all questions together after a longer time interval. Importantly, and again consistent with our hypothesis, in both cohorts there is a robust correlation between ISC and test scores (Fig. 2c; Intentional: r(25)=0.61, *p*=7.51·10^−4^, Incidental: r(29)=0.58, *p*=5.87·10^−4^).

### Different styles of instructional videos

We found a positive correlation between ISC and test scores for all 5 videos tested. The style of these five videos consisted of either animation (lightbulbs, immune, internet) or showed a hand, drawing figures (stars, birth). We wanted to test whether this effect is robust to other popular styles of informal instructional videos found on popular YouTube channels. To this end we performed an additional experiment on a new cohort of 30 subjects (Experiment 3; 22 females, 8 males, age 18-50, mean=25.73, SD=8.85 years) where we selected 6 different videos in three different styles (two videos per style): a real-life presenter along with animation, a presenter writing on a glass board, and writing-hand with animation (see links to videos in Methods section). Despite the different visual appearance and dynamic, we still find a strong correlation between ISC and test scores for all three styles (Fig. 2d, Animation & Presenter: r(27)=0.53, *p*=3.1·10^−3^), Animation & Writing hand: r(28)=0.51, *p*=3.7·10^−3^), Glassboard & Presenter: r(27)=0.46, *p*=0.01).

### Recognition and comprehension questions

It is possible that attention favors recognition of factual information, but that questions probing for comprehension of the material would require the student to disengage from the video to process the content “offline”. We therefore included in Experiment 3 comprehension questions (32 out of a total of 72 questions across the 6 videos, see questions in Tab. 1). Overall subjects did similarly on the comprehension questions as compared to the recognition questions (Fig. 2e) and we find a significant correlation with ISC for these comprehension questions (r(28)=0.50, *p*=5.3·10^−3^), and we again find a correlation with recognition performance (r(28)=0.58, *p*=8.5·10^−4^). These correlation values do not differ significantly (asymptotic z-test after Fisher r-to-z conversion, *p*=0.68) suggesting that comprehension and recognition are both affected by attention. Indeed, test scores for comprehension and recognition questions are significantly correlated across subjects (r(28)=0.52 (*p*=3.02·10^−3^)). Therefore, the hypothesized link between ISC and performance seems to be fairly robust, applying to different learning scenarios, various styles of educational video found online, as well as recognition and comprehension questions alike.

### Capturing eye movements online at scale using standard web cameras

Thus far all experiments were performed in a laboratory setting with a research-grade eye-tracker. However, we showed that the attentional processes for the ISC of both eye movements and pupil size operate at frequencies below 10Hz. This would suggest that moving this technology to more realistic settings using low-grade eye tracking hardware would possible. To test the approach in a realistic setting we developed an online platform that can operate on a large scale of users. The platform relies on standard web cameras and existing eye tracking software that can run on any web browser [23]. The software operates on the remote computer of the users and captures gaze position. In one experiment we recruited 82 students (female=21, age 18-40, mean=19.6, SD=2.7 years) from a college physics class to participate after their lab sessions using the desktop computers available in the classroom (Experiment 4: Classroom). In another experiment we recruited 1012 participants (female=443, age 18-64, mean=28.1, SD=8.4 years) on MTurk and Prolific. These are online platforms that assign tasks to anonymous subjects and compensate them for their work (Experiment 5: At-home). The subjects used the webcam on their own computers emulating the at-home setting typical for online learning. The gaze position data collected with the web camera is significantly noisier than using the professional eye-tracker in the lab (Fig. 3a, see raw data in Supplement S5). To quantify this, we compute the accuracy of gaze position when subjects are asked to look at a dot on the screen (Fig. 3b). As expected, we find a significant difference in gaze position accuracy between the laboratory and the classroom (two-sample t-test t(69)=-7.73, *p*=6.3·10^−11^) and a significant difference between the classroom and the at-home setting (t(242)=-2.46, *p*=0.01). Despite this signal degradation we find a high correlation between the median gaze position data (across subjects) for laboratory and classroom data (Horizontal gaze: r=0.87 ± 0.04; Vertical gaze: r=0.75 ± 0.04) and laboratory and at-home (Horizontal gaze: r=0.91 ± 0.04; Vertical gaze: r=0.83 ± 0.04).

**Figure 3:**
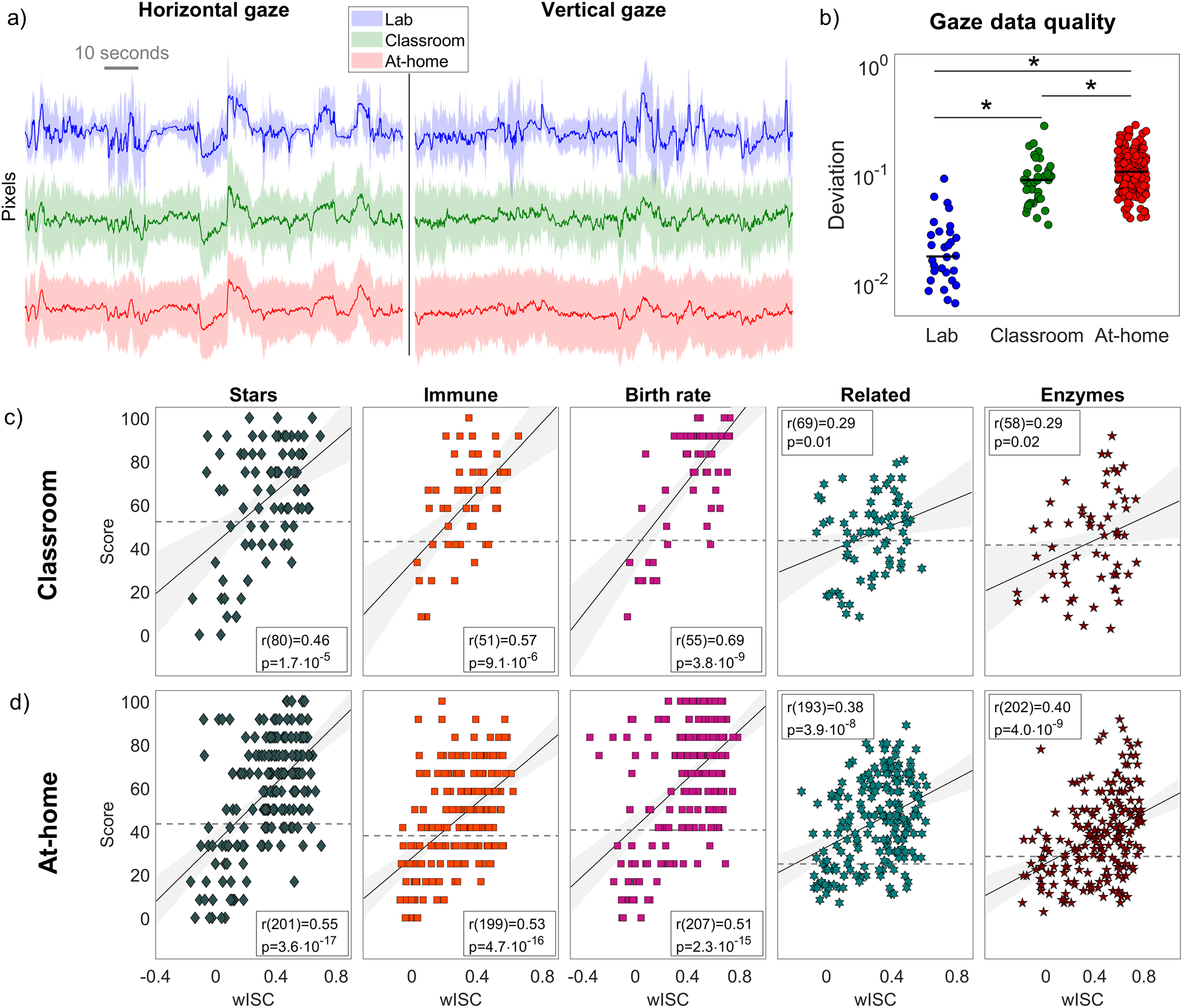
Weighted inter-subject correlation of eye movement measured using low-cost web camera predicts test scores. **a)** Gaze position for ‘Immune’ video in Laboratory, Classroom and At-home settings. Median and interquartile range are taken across subjects (solid line and grayed area respectively). **b)** Deviation of gaze position when subjects looked at 4 “validation” dots presented in sequence on the corners of the screen, collected in the Laboratory, Classroom and At-home settings for the first video shown to subjects (see Methods). *indicates a significant difference in means. **c)** Eye-movement wISC is predictive of performance in the Classroom. **d)** Eye-movement wISC is predictive of performance in the At-home setting.

### Predicting test scores in a classroom and at home using web cameras

To preserve online privacy of the users we propose to evaluate eye movements remotely by correlating each subject’s eye movements with the median gaze positions (Fig. 3a). Instead of ISC with all members of the group, we thus compute the correlation with the median position locally, without the need to transmit individual eye position data (see Methods). To compensate for the loss of the pupil signal we now also measure the correlation of eye movement velocity, which is high when subjects move their gaze in the same direction, regardless of absolute gaze position (see Methods). We combine these eye movement metrics by taking a weighted average of the vertical, horizontal and velocity ISC (wISC; see Methods). We find that this wISC of eye-movement robustly correlates with subsequent test scores (Fig. 3; Tab. S1) despite the lower quality of the gaze position data. In fact, the correlation of wISC with test scores for the classroom (Fig. 3c; r=0.46 ± 0.16, p<0.01) are comparable to the values in the laboratory experiments (r = 0.59 ± 0.08, all p<0.01; compare to Fig. 2b). The at-home experiment had also highly significant correlation between wISC and subsequent test scores (Fig. 3d; r=0.47 ± 0.08, *p*<3.9·10^−8^). The prediction error of the test score is 14.59% ± 16.86% (median across videos, IQR across all videos and subjects), which is equivalent to 1.75 out of 12 questions. We can essentially predict how well a student is going to perform on a test by comparing their eye movements to the median eye movements.

## Discussion

We found that eye movements during watching of instructional videos are similar between students, in particular if they are paying attention. The effect of attention is strong, allowing one to detect with a few minutes of gaze-position data if the student is distracted. Consequently, and as predicted, we find that students performed well in subsequent quizzes if their eyes followed the material presented during the video in a stereotypical pattern. We replicated this finding in two subsequent laboratory experiments, where we confirmed that the effect persists when students do not expect to be quizzed, and that the effect of attention does not depend on the specific type of video or the type of questions asked. The results also replicate in a classroom setting and in a large scale online experiment with users at home using standard web cameras. By correlating with the median gaze-positions one can avoid transmitting personal data over the internet. Thus we conclude that one can detect students’ attentional engagement during online education with readily available technology. In fact, we can predict how well a student will perform on a test related to an instructional video, by looking at their eye movements while maintaining online privacy.

Note that the link between eye-movements and test performance established here was purely correlational. It is possible that stronger students can both follow the video better and also perform better in the test, without the need for a direct link between the two. It could also be that students with prior knowledge on the material were more interested, and thus paid more attention. We built an analytic model assuming a common cause for intersubject correlation and test-taking performance. While we refer to this common cause as “attention”, it really can refer to any internal state of a subject that may have a causal effect on test scores and eye movements such as alertness, interest, motivation, engagement, fatigue, etc. This causal model explained the data more accurately than a simple correlation. But ultimately, our study did not control attention prospectively and thus cannot conclusively answer the direction of this relationship.

We tested for recognition of factual information presented in the videos. Performance on these questions naturally depend on attention to the presentation of this factual information. For questions requiring comprehension, instead, it may be that students need time to think about material quietly without being absorbed by the video. Yet, we did not find a degradation in the ability to predict test scores from eye movement for the comprehension questions. However, a more nuanced analysis and larger sample size may be needed to establish a difference in our ability to predict comprehension vs. recognition performance.

ISC of eye movements varied significantly between subjects and videos. The variability between subjects is to a certain degree predictive of different test scores and thus we can ascribe it to genuine differences in attention. However, there is a significant variability in ISC across subjects even in the distracted condition, suggesting that baseline levels of ISC do vary between subjects, irrespective of attention. ISC also differed significantly among videos. This again could be due to different levels of attention that the videos elicit, but it could also be due to differences in how visual dynamic drives bottom-up attention, e.g. slower videos or less salient visuals may drive eye movements less vigorously [13, 18]. In fact, we occasionally observed short segments of asynchronous eye movements, as gaze jumps back and forth between two prominent items on the video, but out of phase between subjects. Nevertheless, for the full videos ISC was always positive and correlated with learning gains. One caveat of using ISC is that its values may depend on spatial arrangements and temporal dynamic. Therefore, ISC values should always be compared against a baseline that calibrates for differences between videos [24].

Eye tracking has been used extensively to study cognitive processing during reading [25] and visual search [26]. In education research, specifically, eye tracking is often combined with a think-aloud protocol [27], whereby subject verbalize their thinking while learning [28]. This has been used to establish theories of learning [29]. Eye tracking data has also been used to classify different cognitive processes, or to classify whether the viewer is an expert or naive learner [30]. But to our knowledge, eye movements have not been used to predict test-taking performance as we have done here.

Eye tracking has also been used in research specifically related to computer and online education. For instance, when learning from pictures and written text, fixation times and rereading predict learning performance [31]. Showing the instructor’s face while talking seems to help students’ attend to the material [32, 33], but there are mixed results on whether this is actually beneficial for learning [34, 35]. These types of results required careful analysis of the exact content that is fixated upon. The method presented here assesses whether students are paying attention without the need for specific information about the contents of the video.

Given the link between the point of gaze and attention, attempts have been made to guide the attention of novice learners using cueing e.g. pointing out where on a video or animation the student should pay attention. This can reduce cognitive load [36], and foster learning [37]. There are also a few attempts of guiding attention of a novice by displaying the gaze position of an expert during problem solving [38] or during video presentation [39]. However, this method has shown improved learning only in specific cases and requires careful manual annotations of the instructional videos [8].

Our analysis also included pupillary responses. That this should correlate between subjects is perhaps not surprising as it is strongly driven by luminance changes in the visual stimulus. The novel observation is that this correlation is modulated by attention. Interestingly, the ISC of pupil size remained predictive of test taking performance even after regressing out luminance. Therefore, in the present study synchronization of pupil response is unlikely to result from luminance fluctuations and may be driven instead by other factors known to affect pupil size, such as arousal [33], cognitive effort [40] or attention [32]. The present finding differs from the extensive literature on pupil size, which attempt to link pupillary response to specific events. For example, pupil size predicted reliably which stimuli were recalled, in particular for emotionally arousing stimuli [41]. Pupil size has also been linked with cognitive effort, for instance, the effort associated with holding multiple items in working memory [40]. In contrast to this traditional work on event-related pupil dilation we did not have to analyze the specific content of the stimulus. As with the eye movements, we can simply use other viewers as a reference to determine if the pupil size is correlated, and if it is, anticipate high test scores.

Online education often struggles to persistently engage students’ attention, which may be one of the causes for low retention [42]. Student online engagement is often measured in terms of the time spent watching videos [43], mouse clicks [44] or viewed content [45], and some important lessons have been gained from these outcome measures. For instance, videos should be short, dynamic, and show the face of the instructor talking with enthusiasm [43]. Our recent work has focused on measuring attentional engagement of the students by measuring their actual brain activity [46]. Attempts to record brain signals in a classroom have been made [47–49], but typically require help from research personnel and may thus not be practical, particularly at home. The method we have presented here opens up the possibility to measure not just time spent with the material, but the actual engagement of the student’s mind with the material, regardless of where they are. With adequate data quality one may be able to even adapt the content in real time to the current attentional state of the student. In particular, for synchronous online course, where students participate at the same time, real-time feedback to the teachers may allow them to adapt to students’ level of attention in real-time, much like real teachers in real classrooms. The internet has turned attention into a commodity. With video content increasing online, remote sensing of attention to video at scale may have applications beyond education, including entertainment, advertising, or politics. The applications are limitless.

## Methods

### Participants

1182 subjects participated in one of five different experimental conditions. The first two experiments tested the learning scenario of online education, namely intentional learning (Experiment 1, N=27, 17 females, age 18-53 M=26.74, SD=8.98, 1 subject was removed due to bad data quality) and incidental learning (Experiment 2, N=31, 20 females, age range 18-50, mean 26.20, SD 8.30 years; 3 subjects were removed due to bad signal quality). Experiment 3, was designed to investigate the effect of different video styles and types of questions (N=31, 22 females, age 18-50, M=25.73, SD=8.85 years; 2 subjects were removed due to bad signal quality). Participants for the laboratory Experiments 1-3 were recruited from mailing lists of students at the City College of New York and local newspapers ads (to ensure a diverse subject sample). Experiment 4 was designed to replicate the findings from the laboratory in a classroom setting. Participants were all enrolled from a single physics class at the City College of New York (N=82, female=21, age 18-40, M=19.6, SD=2.7 years). Experiment 5 replicated the finding from the laboratory in a home setting. Amazon Mechanical Turk and Prolific was used to recruit subjects (N=1012, 473 female, age range 18-64, M=28.1, SD=8.4 years). Subjects of Experiments 1-4 only participated in a single experiment, i.e. they were excluded from subsequent Experiments. In Experiment 5 subjects were allowed to participate in more than one assignment (video) so the total subject count are not unique subjects. The experimental protocol was approved by the Institutional Review Boards of the City University of New York. Documented informed consent was obtained from all subjects for laboratory experiments. Internet-based informed consent was given by subjects that were recruited for the online experiments.

### Stimuli

The video stimuli are listed in Tab. 1. Briefly, for Experiments 1, 2, 4 and 5 we selected five videos from YouTube channels that post short informal instructional videos: ‘Kurzgesagt – In a Nutshell’ and ‘minute physics’. The videos cover topics relating to physics, biology, and computer science. Videos are short to match the limited attention span online (Range: 2.4 – 6.5 minutes, Average: 4.1 ± 2.0 minutes). Two of the videos (‘Immune’ and ‘Internet’) used purely animations, where ‘Boys’ used paper cutouts and handwriting. ‘Bulbs’ and ‘Stars’ showed a hand drawing illustrations aiding the narrative. For Experiment 3-5 were selected six video stimuli used in using the following criteria: 1. The duration was limited to no more than 6 minutes [43] to ensure our subject would not lose interest (Table 1, Duration: 4.2 – 6 minutes long, Average: 5.15 ± 57 seconds). 2. The videos cover 3 different styles that are commonly found in large online educational channels on YouTube (‘Khan Academy’, ‘eHow’, ‘Its ok to be smart’ and ‘SciShow’). Here we use a nomenclature of video styles as found in the online instructional video community. ‘Mosquitoes’ and ‘Related’ were produced in the ‘Presenter & Animation’ style, which shows a presenter talking as pictures and animations are shown. ‘Planets’ and ‘Enzymes’ were produced in the ‘Presenter & Glass Board’, which shows a presenter drawing illustrations and equations on a glass board facing the viewer. ‘Capacitors’ and ‘Work energy’ used the ‘Animation & Writing hand’ style, which shows a hand drawing animations. 3. Videos within each ‘style’ covers two different topics each related to biology, astronomy or physics. Links to all videos are provided in Tab. 1.

### Procedure

#### Laboratory experiments

In Experiment 1 (intentional learning), subjects watched a video and answered afterwards a short four-alternative forced-choice questionnaire. The 5 videos and question pairs were presented in random order. The subjects were aware that they would be tested on the material. The test covered factual information imparted during the video (11–12 recognition questions per video). Examples of questions and answer options can be found in Tab. 1 and all can be found here. In Experiment 2 (incidental learning) subjects were not aware that they would be tested or asked questions regarding the material. They first watched all 5 videos, and subsequently answered all the questions (59 questions in total). In Experiment 3, subjects were informed that questions regarding the material would be presented after each video and followed the procedure of Experiment 1, using a different set of stimuli with 6 videos. The order of video presentation, questions and answer options were randomized for all three experiments. Common for Experiments 1-3, after subjects had watched all video stimuli and answered questions, they watched all the videos again in a distracted condition using the same order as the attend condition. In this condition participants counted backwards silently in the mind, from a randomly chosen prime number between 800 and 1000, in decrements of 7. This task aimed to distract the subjects from the stimulus without requiring overt responses and is based on the serial subtraction task used to assess mental capacity and has previously been used to assess attention [7].

#### Online experiments

The web camera experiments (Experiments 4 and 5) were carried out using Elicit, a framework developed for online experiments. In Experiment 4 (classroom) students used the same computers they use for their class exercises. From the Elicit webpage subjects could select which video they wanted to watch from a list of 5 videos. Subjects were given a short verbal instruction besides the written instructions that were provided through the website. In Experiment 5 (at-home) subjects could select HITs (Amazon Mechanical Turk assignments) or assignments (Prolific) that contained a single video with questions and otherwise followed the same procedure as Experiment 4. For both Experiment 4 and 5, subjects were informed that there would be questions regarding the material after the video. They first received instructions regarding the procedure, performed the webcam calibration to enable tracking of their eye movements, watched a single video and answered a four-alternative choice questionnaire for that video. Subjects were allowed to perform more than one assignment, i.e. view more than one video and answer questions. In Experiment 5 subjects were additionally shown a short instructional video on how to calibrate the webcam to track eye movements.

#### Online eye tracking using web cameras

The webcam-based gaze position data was recorded using WebGazer [23]. WebGazer runs locally on the subject’s computer and uses their webcam to compute their gaze position. The script fits a wireframe to the subject’s face and captures images of their eyes to compute where on the screen they are looking. Only the gaze position and the coordinates of the eye images used for the eye position computation were transmitted from the subject’s computer to our web server. In order for the model to compute where on the screen the participant is looking, a standard 9-point calibration scheme was used. Subject had to achieve a 70% accuracy to proceed in the experiment. Note that here we did transfer user data to the server for analysis. However, in a fully local implementation of the approach no user data would be transmitted. Instead, median eye positions of a previously recorded group would be transmitted to the remote location and median-to-subject correlation could be computed entirely locally.

### Preprocessing of webcam-based gaze position data

WebGazer estimates point of gaze on the screen as well as the position and size of the eyes on the webcam image. Eye position and size allowed us to estimate the movement of the subject in horizontal and vertical directions. The point of gaze and eye image position & size were upsampled to a uniform 1000Hz, from the variable sampling rate of each remote webcam (typically in the range of 15-100Hz). An inclusion criteria for the study was that the gaze position data should be sampled at least at an average of 15Hz. Missing data were linearly interpolated and the gaze positions were denoised using a 300ms long median filter. Movements of the participant were linearly regressed out of the gaze position data using the estimated head position of the participant from the image patch coordinates. This was done since the estimated gaze position is sensitive to head movements (we found this regression increased the overall ISC). Subjects that had excessive movements were removed from the study (16 out of 1159 subjects; excessive movement is defined as 1000 times the standard deviation of the recorded image patch coordinates in the horizontal, vertical and depth directions). Blinks were detected as peaks in the vertical gaze position data after a 200 ms median filter. The onset and offset of each blink were identified as a minimum point in the first order temporal derivative of the gaze position. Blinks were filled using linear interpolation in both the horizontal and vertical directions. Subjects that had more than 20% of data interpolated using this method were removed from the cohort (14 out of 1159 subjects). We could not compute the visual angle of gaze since no accurate estimate was available for the distance of the subject to the screen. Instead, gaze position is measured in units of pixels, i.e. where on the screen the subject is looking. Since the resolutions of computer screens varies across subjects, the recorded gaze position data in pixels were normalized to the width and height of the window the video was played in (between 0 and 1 indicating the edges of the video player). Events indicating end of the video stimuli (“stop event”) were used to segment the gaze position data. The start time for each subject was estimated as the difference between the stop event and the actual duration of the video. This was done, since the time to load the YouTube player was variable across user platforms.

### Estimating the quality of gaze position

To compute the quality of the gaze position data, subjects were instructed to look at a sequence of 4 dots in each corner of the screen, embedded in the video stimuli before and after the video. The actual dot position on the individual screen was computed and compared to the captured eye gaze position of the WebGazer. The deviation was computed as the pooled deviation of the recorded gaze position from the position of the dot, while the subject looked at each dot. Poor data quality is indicated by higher deviation. Furthermore, subjects with low quality calibration were identified by computing the spatial difference of recorded gaze position data of opposing dots in the horizontal and vertical direction when they were looking at the 4 dots. If the difference in recorded gaze position between dot pairs were in average negative, i.e. left/right reversed, the subject was excluded (135 of 1159).

### Preprocessing of laboratory gaze position data

In the laboratory (Experiments 1-3) gaze position data was recorded using an Eyelink 1000 eye tracker (SR Research Ltd. Ottawa, Canada) at a sampling frequency of 500 Hz using a 35mm lense. The subjects were free to move their heads, to ensure comfort (no chin rest). A standard 9-point calibration scheme was used utilizing manual verification. To ensure stable pupil size recordings, the background color of the calibration screen and all instructions presented to the subjects were set to be the average luminance of all the videos presented during the experiment. In between each stimulus presentation a drift-check was performed and tracking was recalibrated if the visual angular error was greater than 2 degrees. Blinks were detected using the SR research blink detection algorithm and remaining peaks were found using a peak picking algorithm. The blink and 100ms before and after were filled with linearly interpolated values.

### Inter-subject correlation and attention analysis of gaze position data

Inter-subject correlation of eye movements was calculated by (1) computing the Pearson’s correlation coefficient between a single subject’s gaze position in the vertical direction with that of all other subjects while they watched a video. (2) obtaining a single ISC value for a subject by averaging the correlation values between that subject and all other subjects (ISC) (3) and then repeating steps 1 and 2 for all subjects, resulting in a single ISC value for each subject. We repeat these three steps for the horizontal eye movements ISC_*horizontal*_ and the pupil size ISC_*pupil*_. To obtain the measure used for the laboratory experiments we averaged the three ISC values which we call ISC=(ISC_*vertical*_+ISC_*horizontal*_+ISC_*pupil*_)/3. The ISC values for the attend and distract conditions, were computed on the data for the two conditions separately. To test whether ISC varies between the attend and distract conditions, a three-way repeated measures ANOVA was used with fixed effect of video and attentional state (attend vs. distract) and random effect of subject. As an additional measure the receiver operating characteristic curve (ROC) was used. Each point on the curve is a single subject. To quantify the overall ability of ISC to discriminate between attend and distract conditions the area under the ROC curve is used (AUC). To test for the effect motivation, ISC was computed for each video in the attend condition and averaged across all videos. Since the distribution was not Gaussian, we tested for a difference in median ISC values with a Wilcoxon rank sum test. To test for the effect of video style on the attentional modulation of ISC we performed a three-way repeated measures ANOVA. The random effect was subject and fixed effects were stimuli, attentional condition and video style.

### Weighted inter-subject correlation of eye movements

For the experiments with the web camera in the classroom and at-home we compute for each time point in the video the median gaze position across all subjects (Fig. 3a). We then compute the Pearson’s correlation coefficient of that median time course with the time course of gaze position of each subject. We refer to this as median-to-subject correlation, MSC_*vertical*_ and MSC_*horizontal*_. Note that in principle this can be computed with the median gaze positions previously collected on a sample group for each video. To compute this remotely without transmitting the gaze data of individual users, one would transmit this median gaze positions to the remote user of the online platform (two values for each time point in the video). MSC can then be computed locally by the remote user. We additionally compute MSC for the velocity of eye movements as follows. First we compute movement velocity by taking the temporal derivative of horizontal and vertical gaze positions using the Hilbert transform. We form two-dimensional spatial vectors of these velocity estimates (combining Hilbert transforms of horizontal and vertical directions). These vectors are normalized to unit length. The median gaze velocity vectors is obtained as the median of the two coordinates across all subjects. The median-to-subject correlation of velocity, MSC_*velocity*_, is then computed as the cosine distance between the velocity vectors of each subject and the median velocity vector, averaged over time. Finally, we combine the three MSC measures to obtain a single weighted inter-subject correlation value for each subject: wISC = *w*_1_MSC_*vertical*_ + *w*_2_MSC_*horizontal*_ + *w*_3_MSC_*velocity*_. The weights *w_i_* are chosen to best predict test scores with the constraint that they must sum up to 1 and that they are all positive. This is done with conventional constrained optimization. The constraints insure that the wISC values are bounded between −1 and 1. To avoid a biased estimate of predictability we optimize these weights for each subject on the gaze/score data leaving out that subject from the optimization, i.e. we use leave-one out cross-validation.

### Frequency-resolved analysis of ISC

We performed a frequency analysis to investigate at which time scale eye movements and pupil size synchronize between subjects. The vertical and horizontal gaze position signal was band-pass filtered using 5th order Butterworth filters with logarithmic spaced center frequencies with a bandwidth of 0.2 of the center frequency. The ISC was computed for each subject in each frequency band (Experiment 2 on all 5 videos). To obtain a single ISC value per frequency band we averages ISC values for all videos, subjects and across the two directions (horizontal and vertical). The gray-shaded intervals around the mean values (Fig. 1d) are the standard error across subjects. The same analysis was done on the pupil size.

### Student learning assessment

Four-alternative, forced-choice questions were used to assess the performance of students (Score). Test performance was calculated as the percentage correct responses each student gave for each video. For questions that had multiple correct options, points were given per correct selected options and subtracted per incorrect selected option. The questionnaires were designed in pilot experiments to yield an even distribution of answer options from subjects that had not seen the videos. All questions and answer options can be found here.

To estimate the baseline difficulty of the questions, separate naïve cohorts of subjects were given the same questions without seeing the videos. Two different cohorts were recruited from the City College of New York to compare against the cohorts recruited for Experiments 1-4 (Experiment 1,2 and 4, N=26; Experiment 3, N=15) and a third from Prolific to compare against the at-home experiment cohort (Experiment 5, N=25).

All questions were categorized as either recognition or comprehension questions following Bloom’s taxonomy ([50]), and defined here specifically as: Recognition – Question can be answered by remembering a word, phrase, or number which was specifically stated in the video and does not require understanding of scientific concepts to answer correctly; Comprehension – Question involves an application or interpretation of ideas presented in the video or identification of concepts developed in the video that likely requires understanding to be able to answer correctly. To decide on the category for each question, the five authors independently rated each question and the majority rating was selected as the final categorization (see ratings here).

### Relating student test performance and ISC

When evaluating the different learning scenarios (incidental and intentional learning) in Experiments 1 and 2, students’ scores and ISC values were averaged across all videos. ISC was compared to student test performance by computing the Pearson’s correlation coefficient between ISC and test performance. Similarly, to test the effect of video style, the ISC and scores for each subject were averages for the videos produced in different styles and correlated using Pearson’s correlation. Testing the connection between ISC and test scores on each individual video, subjects’ scores were compared with the ISC using Pearson’s correlation. To test whether there is a significant difference in correlation between comprehension vs recognition questions and ISC we used the same ISC values and performed a test between correlation values with a shared dependent variable [51]. Testing how well eye-movement ISC can predict the performance of students on tests regarding the material in the online setting, we use leave-one-out cross validation. We estimate the attention model (see Supplement for description) on all subjects leaving out one subject’s ISC values and their corresponding test scores. We then estimate how well ISC predicts the test score on the left out subject. We do this for all subjects and compute the median absolute deviation between the prediction and the actual score. To test if our eye-movement ISC model is statistically better than a naïve model (only predicting the average score), we subtract the prediction errors of the two models and perform a two-sided sign test.

## Data Availability

The datasets generated and analysed during the current study are available from the corresponding author.

## Author contributions

J.M. and L.C.P. designed the study, J.M. collected the data with help from S.U.J. and P.J.G, J.M. analyzed the data, L.C.P. build and implemented the analytic model, J.M. and L.C.P. wrote the manuscript, R.S. and S.U.J. designed the questions for the student assessment.

## Acknowledgements

We would like to thank Iain Bryson from *Freeroaming Solutions* for his invaluable help in developing Elicit, which allowed us to carry out the online experiments. Without him this would not have been possible. We would like to acknowledge the help of James Hedberg for supporting collection of data from the physics classroom cohort. We thank Uri Hasson, Ofer Tchernichowski and Hernan Makse for helpful comments on an early version of this manuscript. We would like to acknowledge National Science Foundation Grant DRL-1660548 for supporting this project.

## Supplement: Synchronized eye movements predict test scores in online video education

**Table S1:**
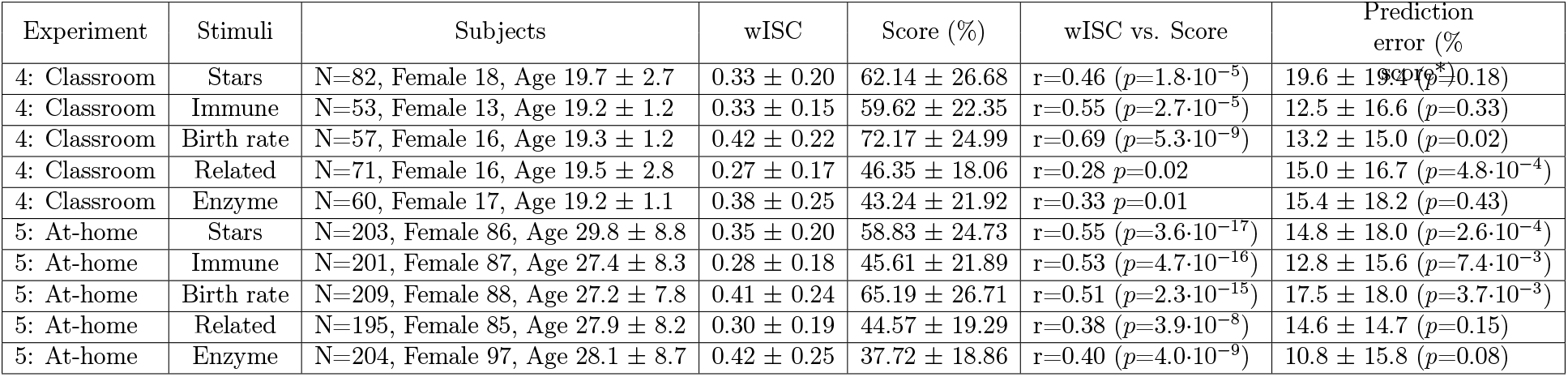
Details of Figure 3. Experiment, stimuli, subjects, weighted Intersubject Correlation (wISC), score, correlation between wISC and score, leave-one-out cross validated Median Absolute Deviation (MAD). Mean ± SD are taken across subjects. *p-values indicate test for whether predicted score is significantly different than naive median score prediction using sign test. z-score is given where this test is approximate.

### S1 Inter-subject correlation of eye movements is modulated by attention

In Fig. 1c) of the main manuscript we showed the result of the attention manipulation task on the ISC of eye movements and pupil size when subjects watched video stimuli. Here we extend this analysis to Experiments 2 and 3.

For Experiment 2 (Incidental learning, Fig. S1a), we perform the same three-way repeated measures ANOVA and find a significant main effect of subject (*F*(29,257)=4.80, *p*=1.65e·10^−12^)), a significant main effect of stimuli (*F*(4,257)=42.08, *p*=4.00e·10^−27^)) and importantly, a significant main effect of attentional state (*F*(1,257) = 1213.27, *p*=2.53·10^−99^)). This suggests that regardless of the different instructions given to the subjects in the two learning scenarios, the measure of ISC is still able to discern the two attentional conditions.

In Experiment 3 six new videos were selected and the attention manipulation task was again used to test the ability of ISC to discern attentional states (Figure S1b). We perform a three-way repeated measures ANOVA and find a significant main effect of subject (*F*(28, 287)=5.41, *p*=1.49·10^−14^), a significant main effect of stimuli (*F*(5, 287) = 19.88, *p*=5.15e·10^−17^)) and importantly, a significant main effect of attentional state (*F*(1,287) = 1357.63, *p*=8.21e·10^−111^)). This indicates the robustness of ISC of eye movements working for a total of eleven different videos.

#### Inter-subject correlation of eye movements modulated by attention for multiple video styles

As education is moving to the online domain, teaching material is growing in abundance and so are the different video styles. We wanted to test if ISC of eye movements as a measure of attentional engagement generalizes to some of the most popular video styles, which are found on the major educational channels of YouTube. In Experiments 1 and 2 subjects watched educational videos produced using the ‘Animations’ (N=3) and ‘Writing hand & Animation’ styles (N=2). In both experiments subjects (N=27,30) watched the videos both in an attentive and distracted condition. The resulting ISC scores for the two conditions can be seen on Fig. S1c).

**Figure S1:**
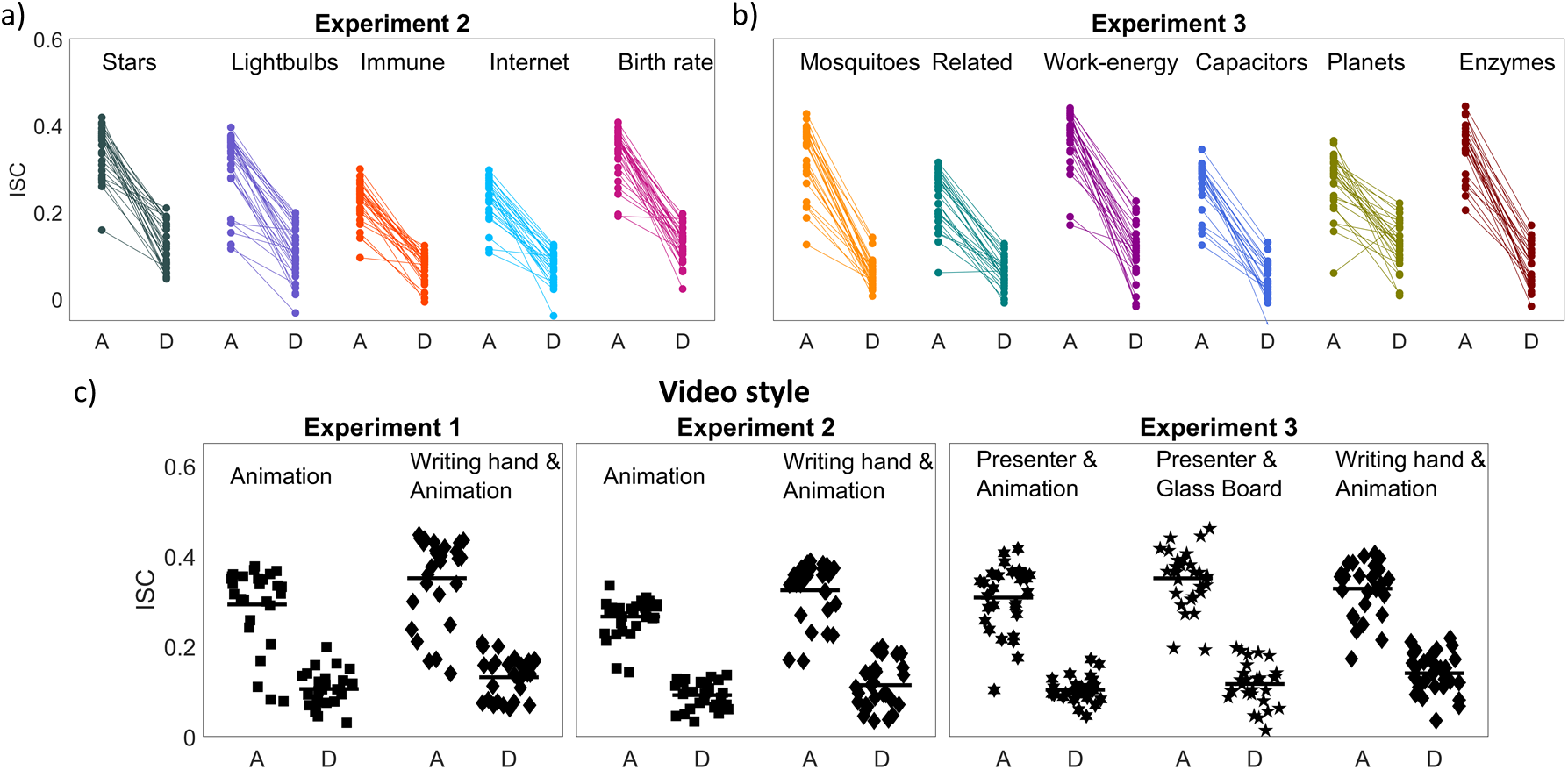
Inter-subject correlation of eye movements and pupil size is modulated by attention when watching educational videos. **a-b)** The intersubject correlation (ISC) values for each subject are shown as dots for all video tests in Experiment 2 (panel a and 3 (panel b). Each dot is connected with a line between two different conditions for the same subject, namely, when subjects were either attending (A) or were distracted (D) while watching the video. **c)** The intersubject correlation (ISC) values for each subject averaged across videos of different styles for Experiments 1-3.

Importantly, in Experiment 1 we find that ISC is modulated by attention for both video styles (Fig. S1c, left), performing a three-way repeated measures ANOVA with subjects as a random effect and attentional state and video style as fixed effects. We find a significant main effect of attentional state (*F*(1,260)=638.60, *p*=5.63·10^−72^)) but no significant main effect of video style (*F*(1,260)=0.43, *p*=0.51).

These finding are replicated in Experiment 2 using a different learning scenario on a new cohort (Fig. S1c, middle). Here we perform the same two-way repeated measures ANOVA test and find a main effect of attentional state (*F*(1,289)=875.74, *p*=1.83·10^−89^)) and no significant effect of video style (*F*(1,289)=0.38, *p*=0.54). This indicates that with this small sample we do not see any effect of video style on the ability of ISC to discriminate between attentional states.

In Experiment 3 we extended our analysis by including 2 additional video styles, namely ‘Presenter & Animation’ and ‘Presenter & Glass Board’ (Figure S1c, right). Despite these very different video styles we again find a robust discrimination of attentional state with the eye movement ISC (*F*(1,287) = 1365.79, *p*=4.03·10^−111^)). However, in this case we do find a main effect of video style (*F*(2,287)=20.57, *p*=4.47·10^−9^)). We attribute this to the general dynamics of the video, where some have high spatial dynamics whereas others are more static eliciting less eye movements. Despite these differences, regardless of video style or learning scenario we find a robust discrimination between attention conditions replicated on three different cohorts.

### S2 Inter-subject correlation of eye movements predicts test scores

In the main text we report a significant correlation between test score and ISC and show the results for Experiment 1 (in Figure 2b). Here we show the same results for Experiments 2 and 3 (Fig. S2a and S2b respectively).

In the main text we analyzed video style in Experiment 3. Here we report similar results for Experiment 1 (Figure S2c, left) for video styles ‘Animation’ (r=0.67, *p*=1.1·10^−4^), N=27) and ‘Writing hand & Animation’ (r=0.62, *p*=5.4·10^−4^), N=27). We reproduce these findings in Experiment 2 using a new cohort and learning scenario (Figure S2c, right). We find a significant correlation for ‘Animation’ (r=0.40, *p*=0.03, N=30) and ‘Writing hand & Animation’ (r=0.50, *p*=5.0·10^−3^), N=30).

### S3 Modeling ISC and test scores as result of measurement noise and attention as common cause

The test scores were determined with a short quiz of only 11-12 questions per video. This makes the scores inherently noisy. Noise in the eye tracking data also seems to have affected the accuracy of the ISC metric. To determine how these noise sources limit our ability to predict test scores we formulated a probabilistic model (Supplement S4). This model aims to match the observed distribution of test scores and ISC (scatter plots in Fig. 2) and 3). The simplest model captures the correlation between scores and ISC assuming they are normally distributed (Fig. S4a, Gaussian model). We also build a model based on the hypothesis that attention causally affects eye movements as well as test scores (Fig. S4b, Attention model). We fit both models using maximum likelihood optimization (see Supplement S4 for details) and find that the causal attention model is significantly more likely than the simple correlation model given the data (Fig. S4c). The parameters of the attention model are consistent with independent empirical observations, such as the baseline performance of naïve subjects (Fig. S4d), or the noise estimates of the eye movements (Fig. S4e). According to the model the differing performance in predicting test scores is well explained by a change in signal-to-noise ratio (SNR, Fig. S4f). SNR is an estimate of the variance in attention over the variance in the noise of the attention measure (eye movement ISC). If SNR could be improved, then prediction performance could be substantially improved (Fig. S4f). However, with only 12 test questions there is a limit to this prediction performance as the test scores are inherently noisy. For instance, despite the relatively low eye tracking noise in the laboratory experiment (both measured and estimated; Fig. S4f) the model suggests that predictability is limited to r<0.7 (Fig. S4g), which is consistent with the empirical data (Fig 2b). Thus, in order to achieve better prediction of test scores one would need to increase the number of questions to obtain a more reliable assessment of student performance (Fig. S4g). In summary, a larger number of test questions, lower noise in our estimate of the attentional state of the subjects, and larger variance in attention across subjects are all expected to contribute to better prediction of test scores.

**Figure S2:**
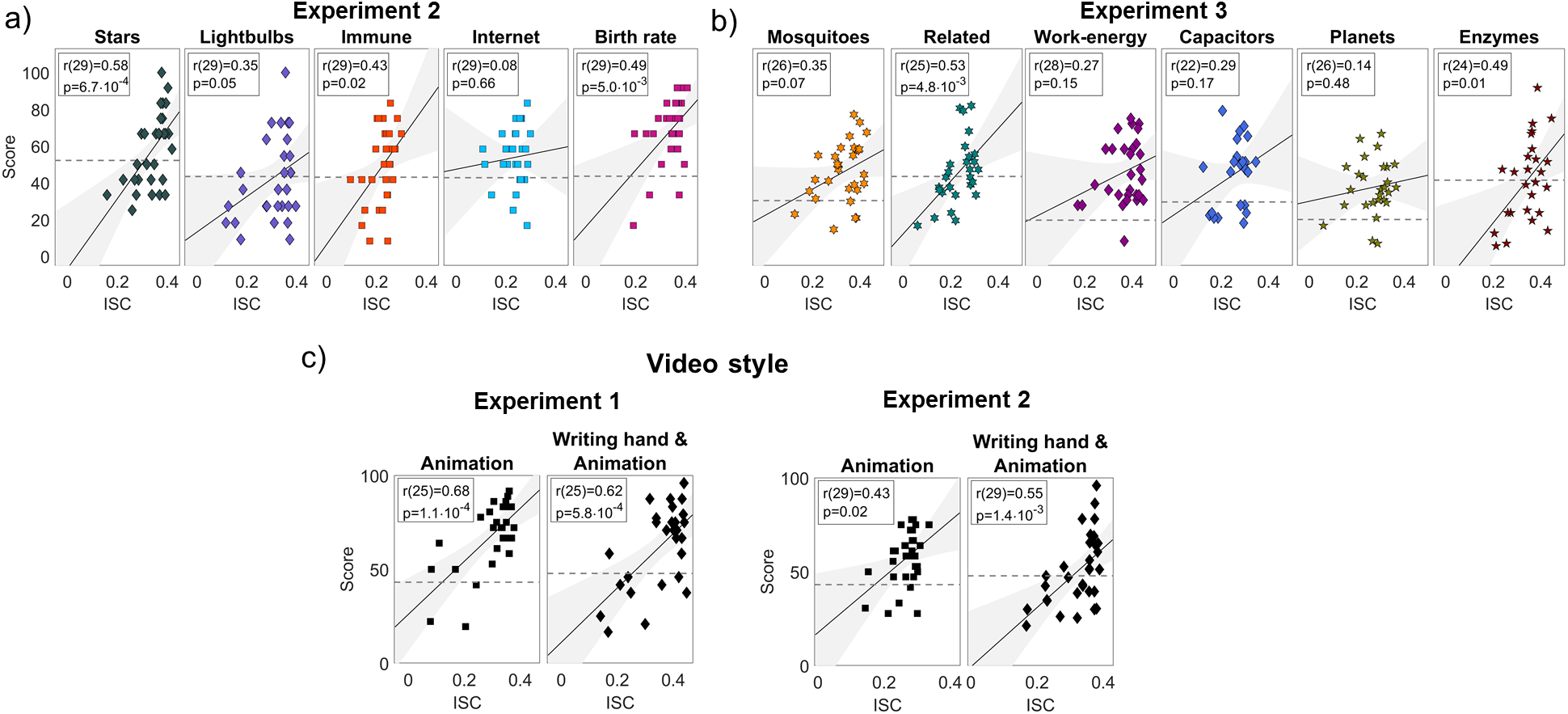
Inter-subject correlation of eye movements predict test scores. **a)** The relation between intersubject correlation of eye movements and pupil size and performance on test taking (Score) for each of five videos of Experiment 2. Each dot is a subject. **b)** same as panel (a) but for the six videos in Experiment 3. **c)** same as panel (a) but averaging over the video produced in the ‘Animation’ and ‘Writing hand & Animation’ styles for Experiments 1 and 2.

### S4 Probabilistic model of relationship between ISC and test scores

#### The data likelihood

The goal of this Supplement is to formulate a probabilistic model for the observed data, namely, the test scores *k_i_* and the correlation values *c_i_* of eye movement. These are measured for subjects *i* =1…*N*. First we assume that these measures are independent across subjects, so that the data likelihood factorizes:

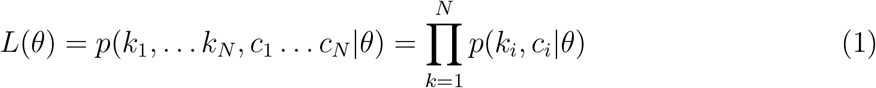

Here *p*(*k, c|θ*) is the joint probability density of the data given parameters *θ*. Next we propose two generative models for this joint density. One is a straightforward bivariate Gaussian density that captures the correlation between the two variables. We refer to this as the “Gaussian model” (Figure S3a). The other will incorporate a common cause that leads to observations *k* and *c* through specific processes. We refer to that as the “Attention model” (Figure S3b).

**Figure S3:**
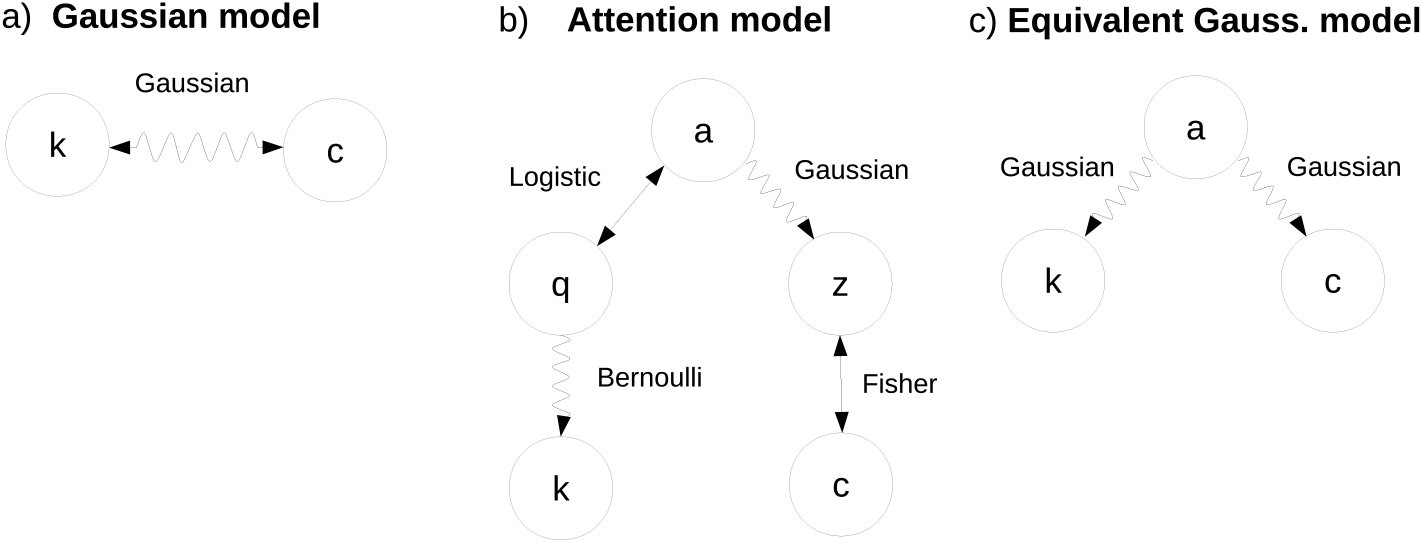
Alternative models to explain test scores *k* and ISC values *c*. Wavy lines indicates probabilistic relationship between variables. Straight line indicates deterministic relationship. Two-sided arrows indicate that the relationship is invertible. **a)** The two variables are related by a bivariate Gaussian distribution. **b)** Variable *a* indicates “attention”, or more generally, an unobserved internal state of the subject that affects test scores and eye movements. Variable *q* captures the odds of answering a question correctly, i.e. questions to the subjects are Bernoulli trials with odds *q*, and the sum of correct answers is test scores *k*. Variable *z* is the Fisher transformed version of correlation value *c*. It is assumed that this variable captures attention, except for additive Gaussian noise. **c)** In the Gassian model the correlation of *k* and *c* can be equivalently formulated as the result of a common driving factor *a* that is also Gaussian distributed.

#### Gaussian model

The canonical approach to describing the correlation between two variables as a probability density is the bivariate Gaussian density

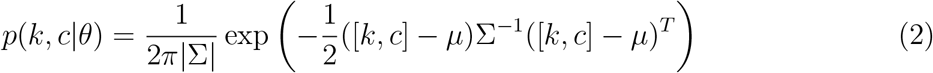

The maximum likelihood solution for the parameters *θ* = {*μ*, Σ} can be found in any statistics text book. They are the sample mean and sample covariance:

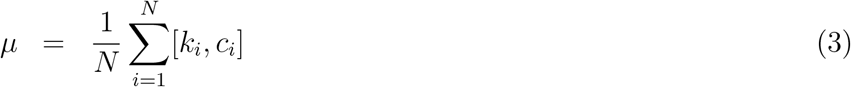

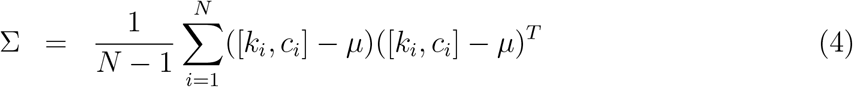

Note that we have here 5 parameter that have been fit to the data: *μ* = [*μ_k_, μ_c_*], and 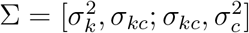.

#### Attention model

The second approach is an explicit formulation of our hypothesis that attention affects both test-taking performance and how similar eye movements are between subjects. We assume that variables do not otherwise affect each other, i.e. eye movement don’t directly affect test scores, and evidently, test scores can not affect eye-movements that happened in the past. In this view, the joint density can be written as

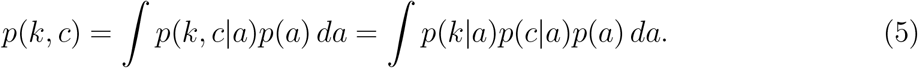

Here variable *a* quantifies the level of attention, and it is distributed across subjects according to *p*(*a*). We do not take the word “attention” here too literally. From a modeling point of view, this variable captures any internal state of the subject that affect performance as well as eye movements. This could include alertness, engagement, interest or fatigue. For instance, a low value for *a* could be due to fatigue or lack of interest in the material. As an internal state of the subject, *a* is an unobserved variable, so we integrate over it. We assume that once conditioned on attention *a*, the score *k* and eye correlation *c* become independent with *p*(*k|a*) representing the probability that at a given attention level a students obtains a score of *k* in the subsequent exam, and *p*(*c|a*) is the probability that at a given attention level the eye movements correlates with other subjects at a value *c*. We will now specify an analytic model for each of these terms.

We assume that attention *a* is normally distributed in our cohort with standard deviation *σ_a_* and zero mean, where zero represents an average level of attention, while positive and negative values represent more or less than average attention.

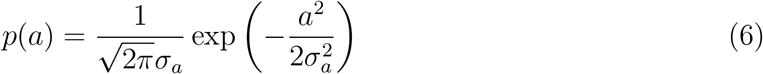

The exam consists of a series of yes/no questions, lets say, *n* questions. Assume that a student has a chance *q* of getting each question right. The number of correct answers *k* is then Bernoulli distributed:

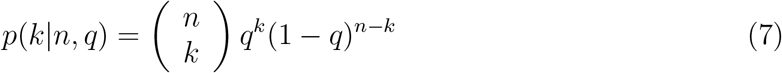

To link attention to performance we assume that high attention levels will give students a high chance of answering questions correctly (close to *q* = 1) and low attention will cause poor odds (close to *q* = 0). Denote with *θ_a_* the level of attention at witch a subject reaches even odds of answering correctly (*q* = 0.5), and let *β_a_* be the sensitivity of performance on changing levels of attention. Then we can represent odds of correctly answering a question as a function of attention as follows:

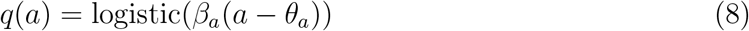

where we used the logistic function to capture the transition from probability 0 to probability 1:

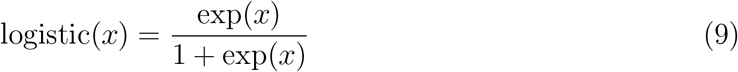

Now we turn our attention to the distribution of correlation values *c*. The density of correlation values is well characterized by a normal distribution after the Fisher z-transformation:

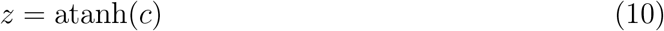

We will therefore work with the density *p*(*z|a*) instead of *p*(*c|a*). The two can be related after appropriate scaling:

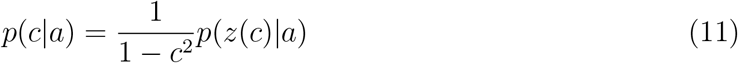

Given our finding that attention strongly modulates the correlation values we assume that *z* is directly determined by attention. However, we have seen that there are different noise levels in measuring eye movements, and so we will assume that *z* carries an independent additive noise

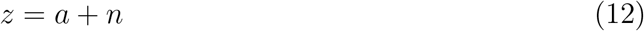

with *n* normally distributed with mean *μ_n_* and standard deviation *σ_n_*. Given this normal noise, *z* given *a* is distributed with the following density

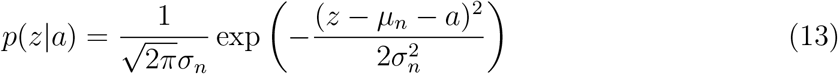

In total, the joint distribution of the model can be written as

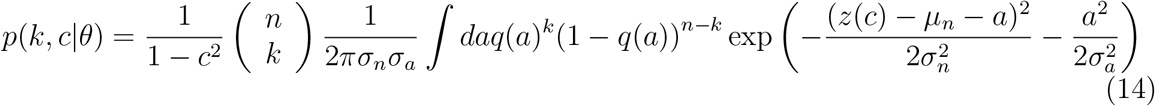

The parameters of this joint density are now *θ* = {*σ_a_, β_a_, θ_a_, σ_n_, μ_n_*,}. To estimate these 5 parameters we will again use maximum likelihood optimization.

For the purpose of finding the optimal parameters first recall that *n* and *a* are independent and that *a* is zero mean. Therefore the following constraints apply to the parameters:

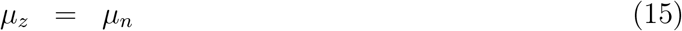

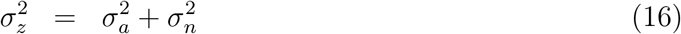

Now, *μ_z_* and 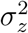 can be estimated directly from the sample data:

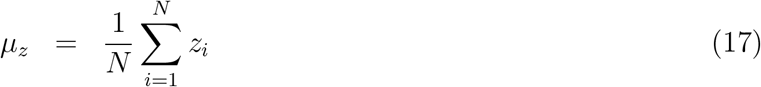

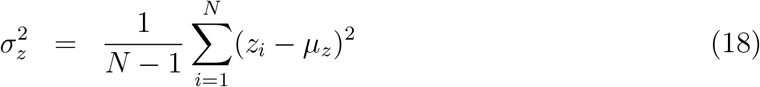

With these two constraints determined by the data, we really only have 3 degrees of freedom remaining for optimization. Parameter *μ_n_* is directly specified by *μ_z_* (15). Parameters *σ_a_* and 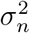 will be reduced to a single parameter, namely, the logarithm of the signal to noise ratio:

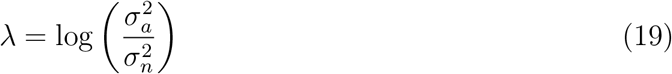

by leveraging the constraint (16) the two variances are quantified with a single parameter λ:

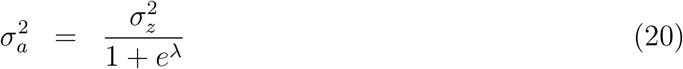

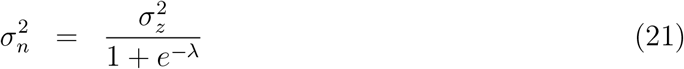

So in total we directly measure *μ_n_* and optimize for the log-likelihood with respect to parameters {λ_*a*_, *β_a_, θ_a_*}. Optimization of three unconstrained parameters can be done fairly efficiently numerically.

Unfortunately the integral over *a* in (14) has no obvious solution and we thus evaluated it numerically during optimization. To do this with equal accuracy for arbitrary parameter values we perform the following variable substitution: *a*′ = *a* – *μ_n_* and write the integral as

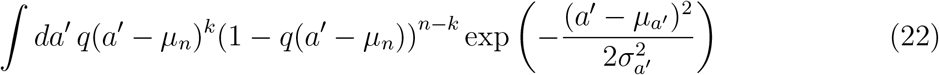

with

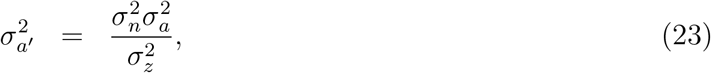

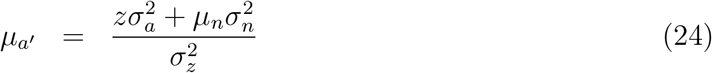

In practice the integral is executed as a sum by evenly dividing the range 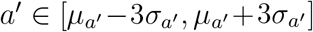 into discrete steps of Δ*a*′ = 0.1*σ_a′_*.

#### Comparison between models

Here we want to make a few concluding remarks in comparing these two models. Both the Gaussian model and the Attention model have 5 free parameters that are fit to the data (Figure S4a and S4b respectively). According to the Akaike Information Criterion one can choose the preferred model by comparing the maximum likelihood value obtained on the data. In fact, since both models have the same number of parameters no correction is needed for the degrees of freedom and one can directly evaluate the likelihood ratios. These ratios are reported in Figure S4c of the main paper for all experimental data. We find that the Attention model is significantly more likely in all cases (likelihood of Attention model over Gaussian model is larger than 1).

The parameters of the two models quantify similar properties of the distribution: *μ_n_* and *μ_c_* take on identical roles, and *σ_c_* is captured by *σ_a_* and *σ_n_*; *θ_a_* affects the mean score *μ_k_* and *β_a_* and *σ_a_* affect *σ_k_*; finally the larger *σ_n_* (more noise), the smaller will be the correlation between *k* and *c*, as captured by *σ_kc_*.

The variables of the Attention model, once fit to the empirically observed data (performance scores and eye movement ISC) seem to correctly capture empirical observations that were assessed independently of this data. For instance, the performance of naive subjects decreases with the estimated attention threshold *θ_a_* (Figure S4d). This is expected as *θ_a_* intends to capture the difficulty of questions (how much attention is required to answer half the questions correctly, in average). Thus, videos with easy tests should have a low estimate for *θ_a_* and videos with harder tests should have a high estimate. Additionally, the measured deviation from fixation dots increases with the estimated noise *σ_n_* (Figure S4e). This is expected as *σ_n_* captures the noise in the eye-movement ISC. Thus, inaccurate eye tracking data should lead to noise in the measured eye-movement ISC. Finally, the estimated signal-to-noise ratio *σ_a_/σ_n_* across videos follows the expected relationship with correlation (of score vs ISC) as predicted by the model (Figure S4f). While these estimates are not independent from the data used for the parameter fit, it does show internal consistency of the modeling approach across videos and supports the argument that the correlations between score and ISC differ due to differing SNR levels of the eye-tracking.

**Figure S4:**
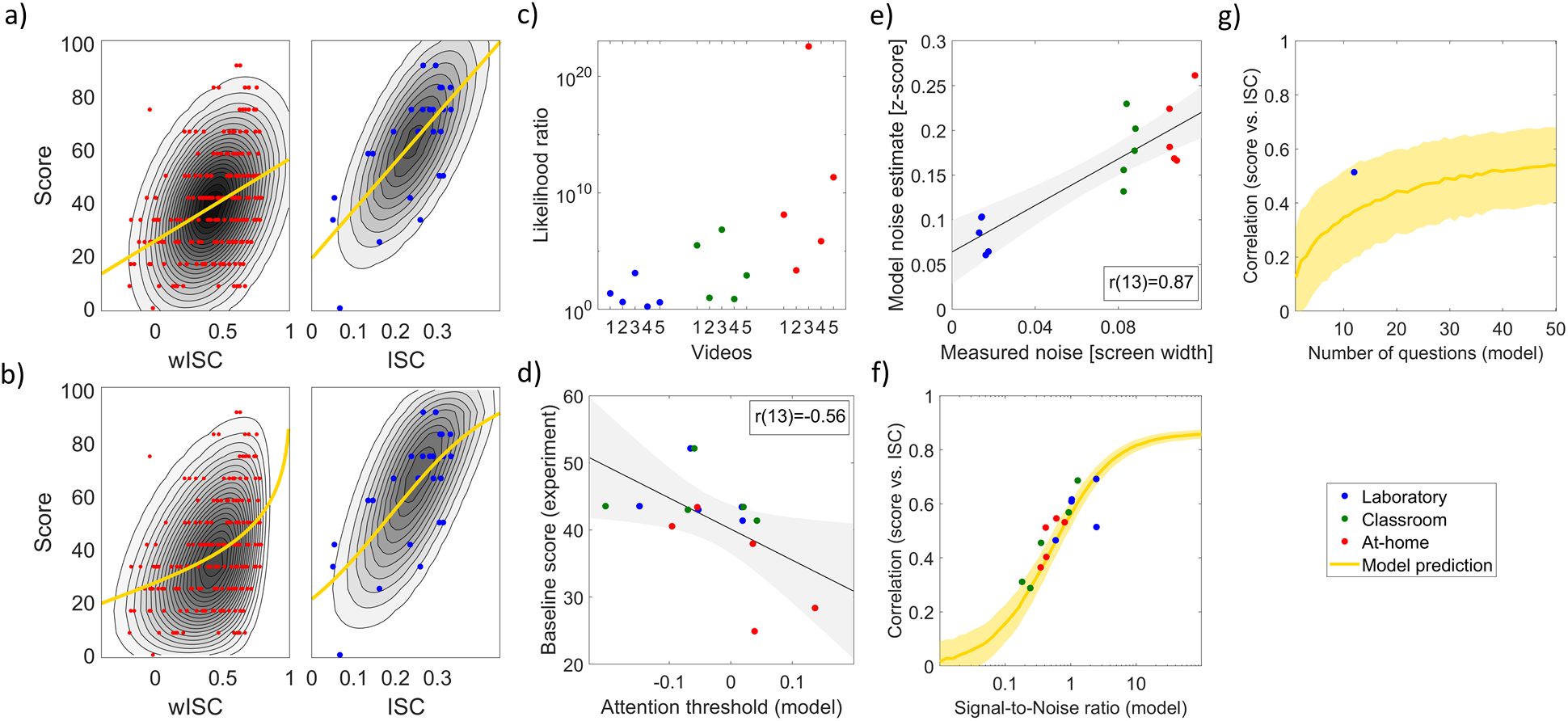
Analytic model of test scores and ISC of eye movements: **a)** Gaussian model fit to laboratory and at-home experiments for videos ‘Immune’ (blue dots) an ‘Birth rate’ (red dots) respectively. This model assumes a bivariate Gaussian distribution. Contour lines indicate the likelihood according to the model. **b)** Fit of the Attention model for the same data. This model assumes that score is Bernoulli distributed and ISC follows a Fisher z-score (see Supplementary Fig. S3). **c)** likelihood ratio between the Attention model and the Gaussian mode for all data from experiments in laboratory (Figure 2b), classroom (Figure 3c), and at-home (Figure 3d). Values larger than 1 indicate that the Attention model is a better fit than the Gaussian model. **d)** Performance of naive subjects, i.e. baseline test scores, compared to the estimated attention threshold in the model *θ_a_*. This threshold parameter is the level of attention required to achieve 50% correct answers. **e)** comparison of estimated noise *σ_n_* and measured noise (Deviation as in Figure 3b). **f)** correlation between eye-ISC and test scores as a function of estimated signal-to-noise ratio, 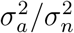. The variability observed in the empirical data is consistent with changing SNR. **g)** correlation between ISC and test scores as a function of the number of questions. Predictability of test scores increases with number of questions. Yellow curve indicate mean model prediction. In panel f and g the model is fit to the data for one of the videos in the laboratory experiment (the one shown in panel a); then data is sampled from this model at random (with SNR or number of questions adjusted according to horizontal axis in panels f and g respectively). Bold curve and shaded area indicate mean and and 95% confidence interval over 1000 random samples.

The Gaussian model can be equivalently described in terms of a common cause (Figure S3c), i.e. the correlation between variables *k* and *c* is introduced by an unobserved common cause *a*. In the case of normally distributed variables the Gaussian model (Figure S3a) and the Equivalent model (Figure S3c) are mathematically indistinguishable. The Gaussian model (Figure S3a) can also be described as normal distributed variable *k* affecting *c* with some Gaussian noise, or vice versa. So one can not discern the direction of causality or a common cause for Gaussian data. For the Attention model the direction of influence can not be readily reversed. Thus, a better fit of the Attention model may be suggestive of a causal relationship via an unobserved common cause.

The Attention model takes the bounded nature of the variables *k* and *c* explicitly into account, whereas the Gaussian model does not. Thus, it is no surprise that the attention model has a higher likelihood on the empirical data as it does not allocate any probability mass outside the valid data range. In contrast, the Gaussian distribution has non-zero probability outside the valid range of the data. So a better fit may simply reflect that the data is more carefully modeled. Indeed, if the Bernoulli distribution is replaced by a Gaussian and the logistic and Fisher transformations by a linear transformation, then the attention model of Figure S3b reduced to the equivalent Gaussian model of Figure S3c. Ultimately the two models are not very different except for this features of a more careful model of the observed variables.

### S5 Gaze position data collected in at-home experiment

To provide a sense of the raw data here we display gaze position collected using web cameras in the at-home Experiment for one of the videos (‘Birth rate’) for all subjects (Figure S5). Each row is a subject and the intensity is the eye gaze position. Subjects are sorted by how well they did on the test that followed the video. It seems clear that high-performing subjects have a stereotypical pattern of eye movements and this pattern disappears as performance drops.

### S6 ISC of pupil and eye movements

In our experiments we have combined the intersubject correlation of eye movements in the horizontal and vertical direction together with the pupil size. Here we take a look at how each of these modalities predict test score independently. First we look at how predictive eye movements are of test taking performance. We average the intersubject correlation of eye movements in the horizontal and vertical directions while 29 subjects watched 5 different instructional videos in Experiment 2. On Fig S6 we see that using eye movements alone we find a significantly correlation with test taking performance (r(27)=0.52, *p*=4.2·10^−3^) and equally with pupil size (r(27)=0.47, *p*=0.01).

**Figure S5:**
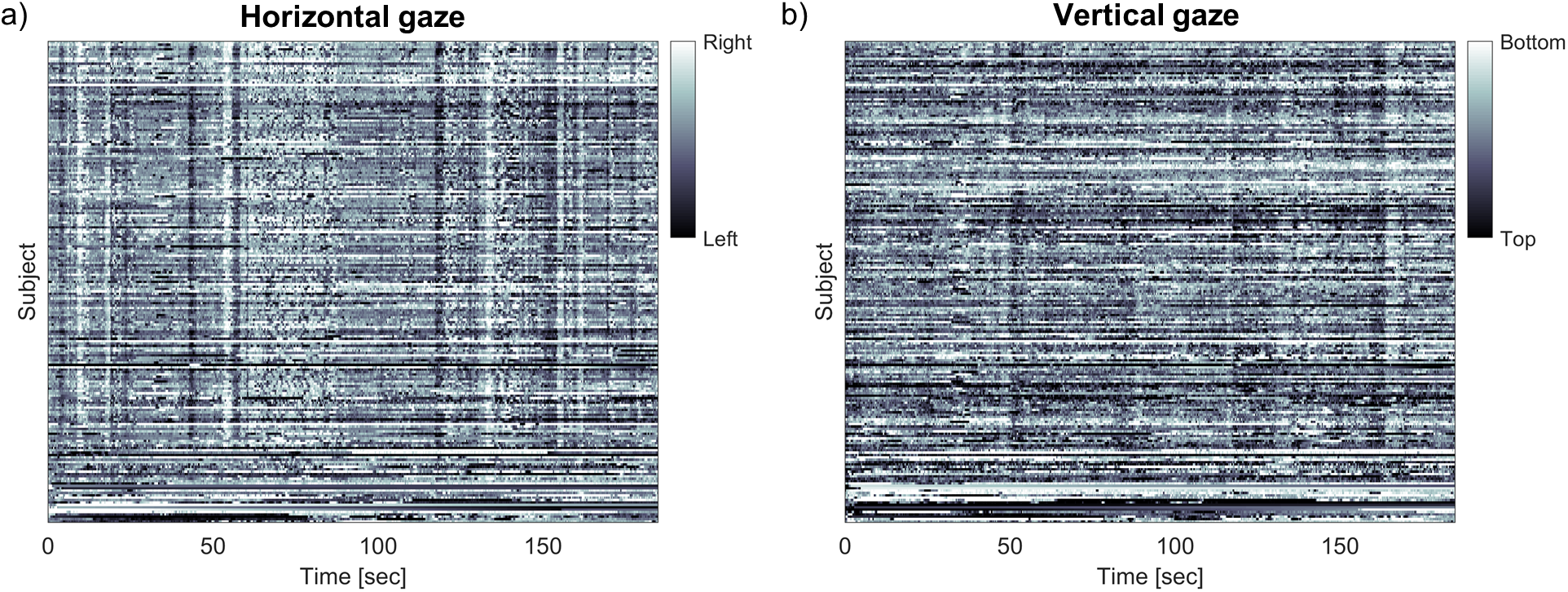
Raw gaze position data: Gaze position collected for ‘Stars’ in the at-home condition. Position is coded as brightness, and each subject is a row. Subject (N=203) are sorted by the score they obtained in the subsequent test (highest on top).

### S7 Removing the effect of luminance on pupil size

The correlation of pupil size across subjects might be driven by the common luminance fluctuations in the video. To test for this we regressed out the luminance from the pupil size for each subject, using both the global luminance (of the entire video frame), as well as the local luminance (5 pixel radius around the current point of gaze, on a 19” monitor with 1280×1024 resolution). For regression we used a time-window of 0-2.4 seconds at 8ms resolution (i.e. 300 samples). Due to the well-known logarithmic relationship between luminance and pupil size [52] we in addition added log transformed pixel luminance values to our regressors. Computing ISC of this pupil signal with local luminance removed we find higher ISC values (M=0.37, SD=0.05) compared to the raw pupil signal itself (M=0.19, SD=0.05, t(28)=3.1193, *p*=4.2·10^−3^, d=0.058). More importantly we find a significant correlation between pupil ISC and test taking performance (r(27)=0.44 *p*=0.02; Fig. S6, center). A similar result is obtained when removing global luminance (Fig. S6, right) (r(27)=0.46, *p*=0.01). Therefore, synchronization of pupil response is unlikely to result from luminance fluctuations and may be driven instead by other factors known to affect pupil size, such as arousal [33], cognitive effort [40] or attention [32] amongst others.

**Figure S6:**
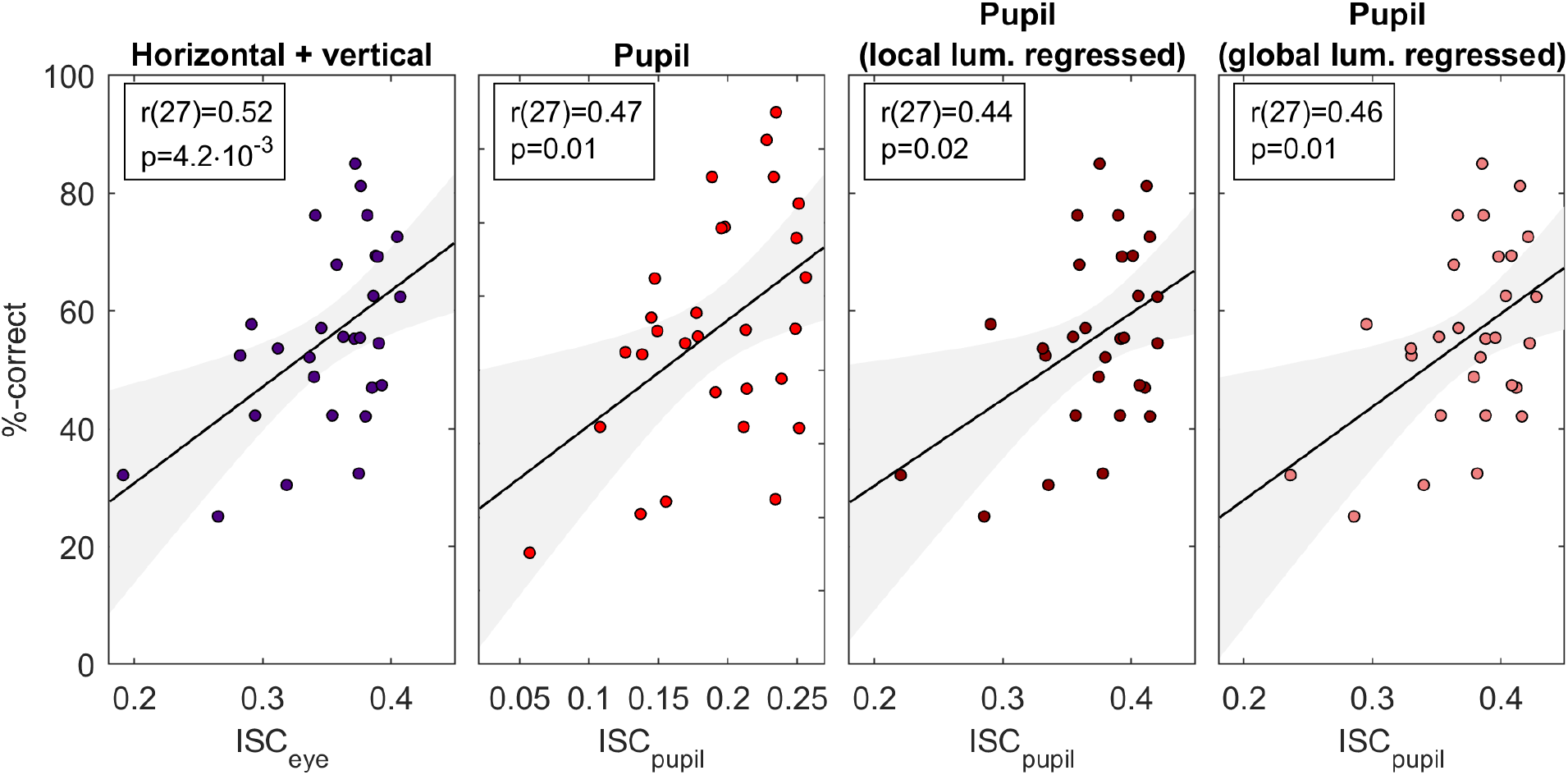
Inter-subject correlation of eye movements and pupil size predict test scores, even when local or global luminance is regressed out.

